## Supplementary figures and images for "Dorsoventral-mediated *Shh* induction is required for axolotl limb regeneration"

### Sup. Fig. 1

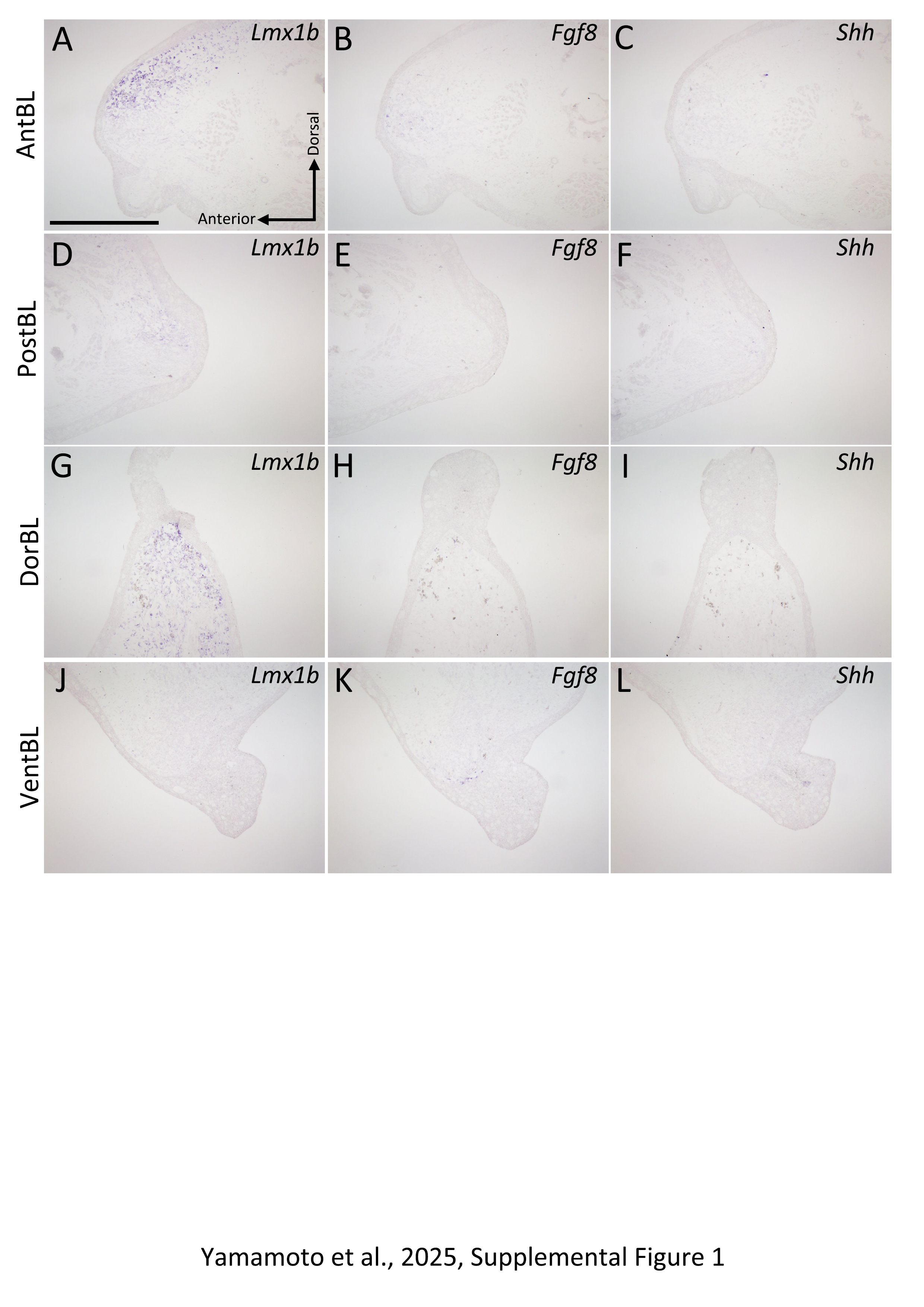

### Sup. Fig. 2

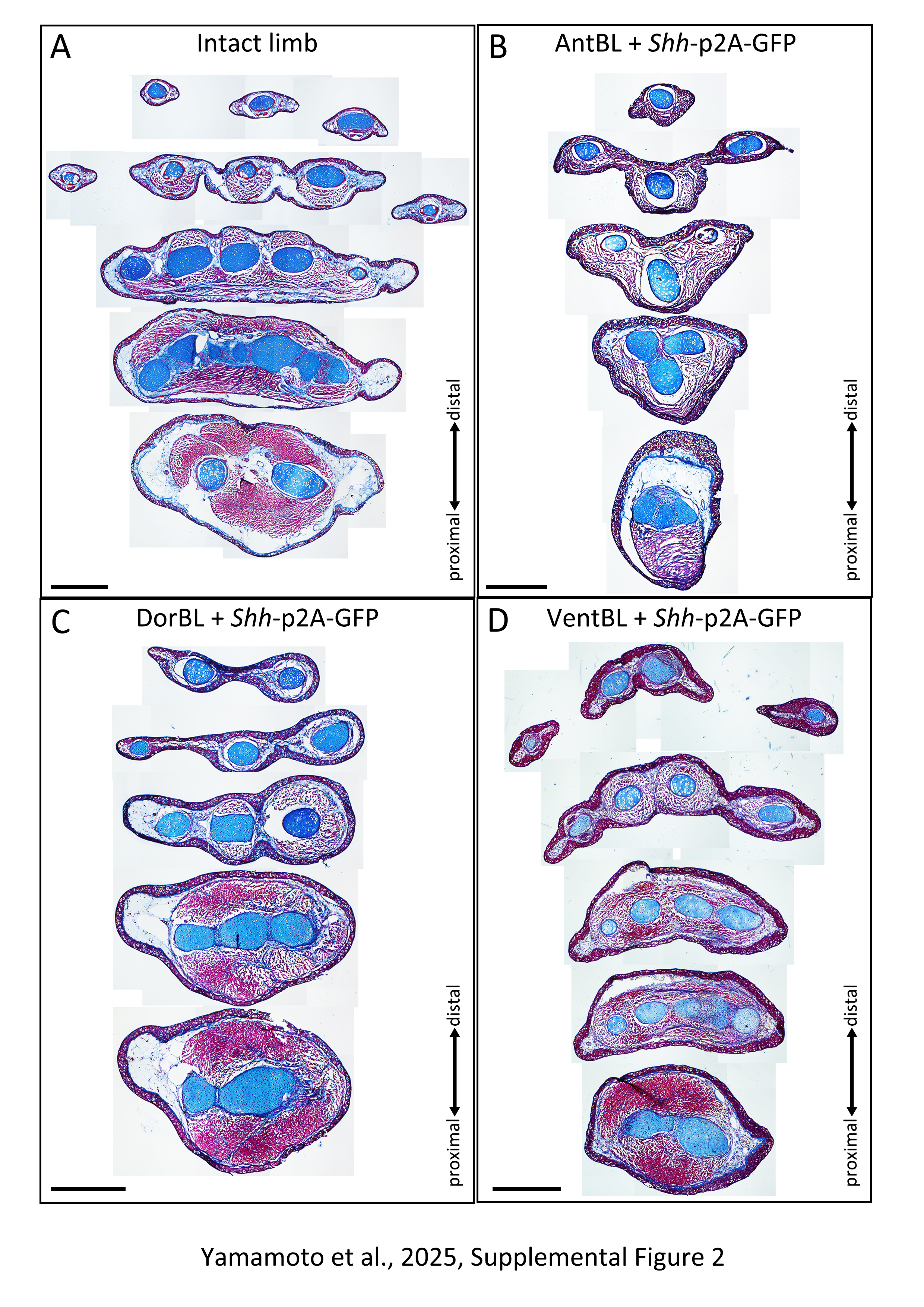

### Sup. Fig. 3

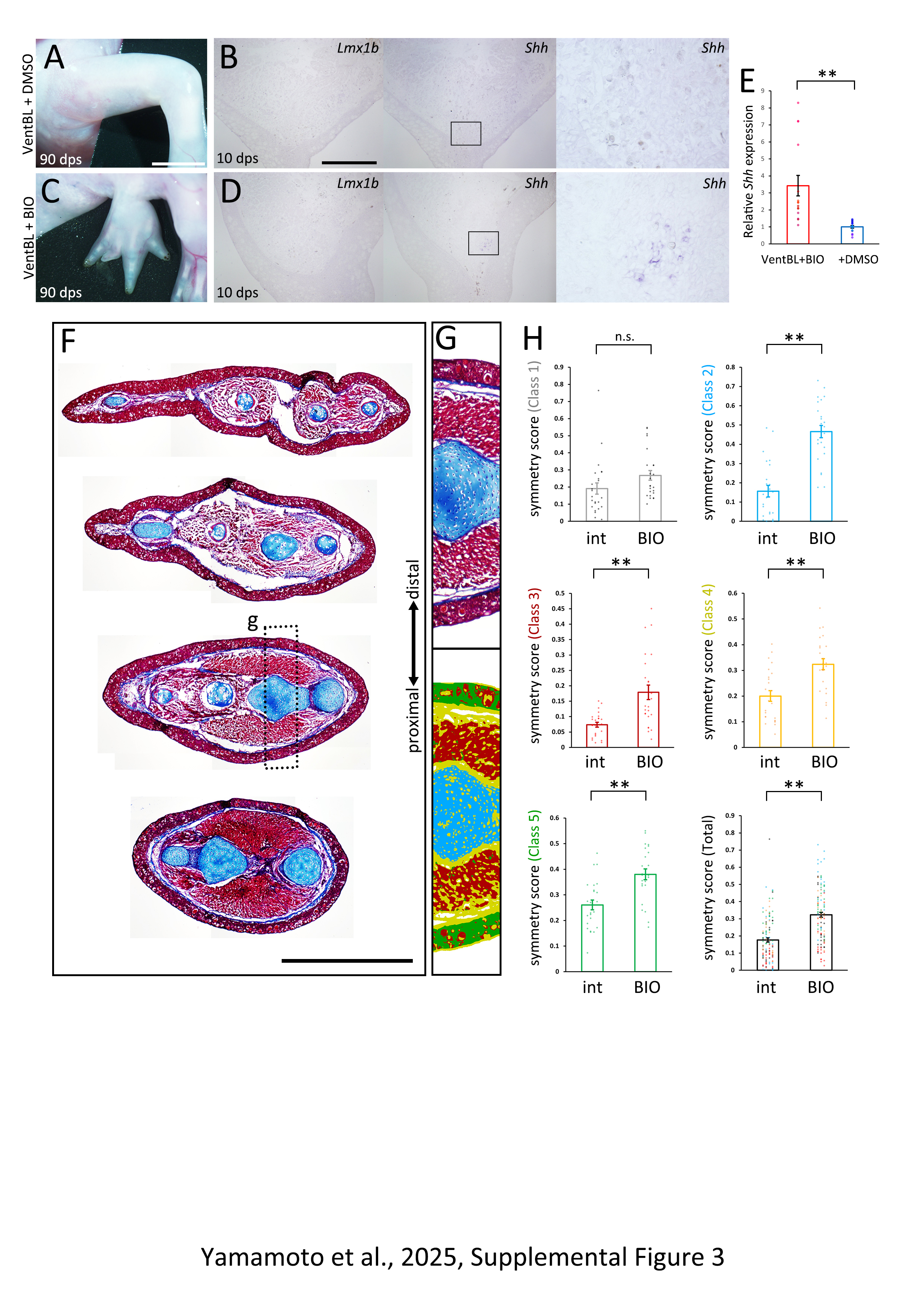

### Sup. Fig. 4

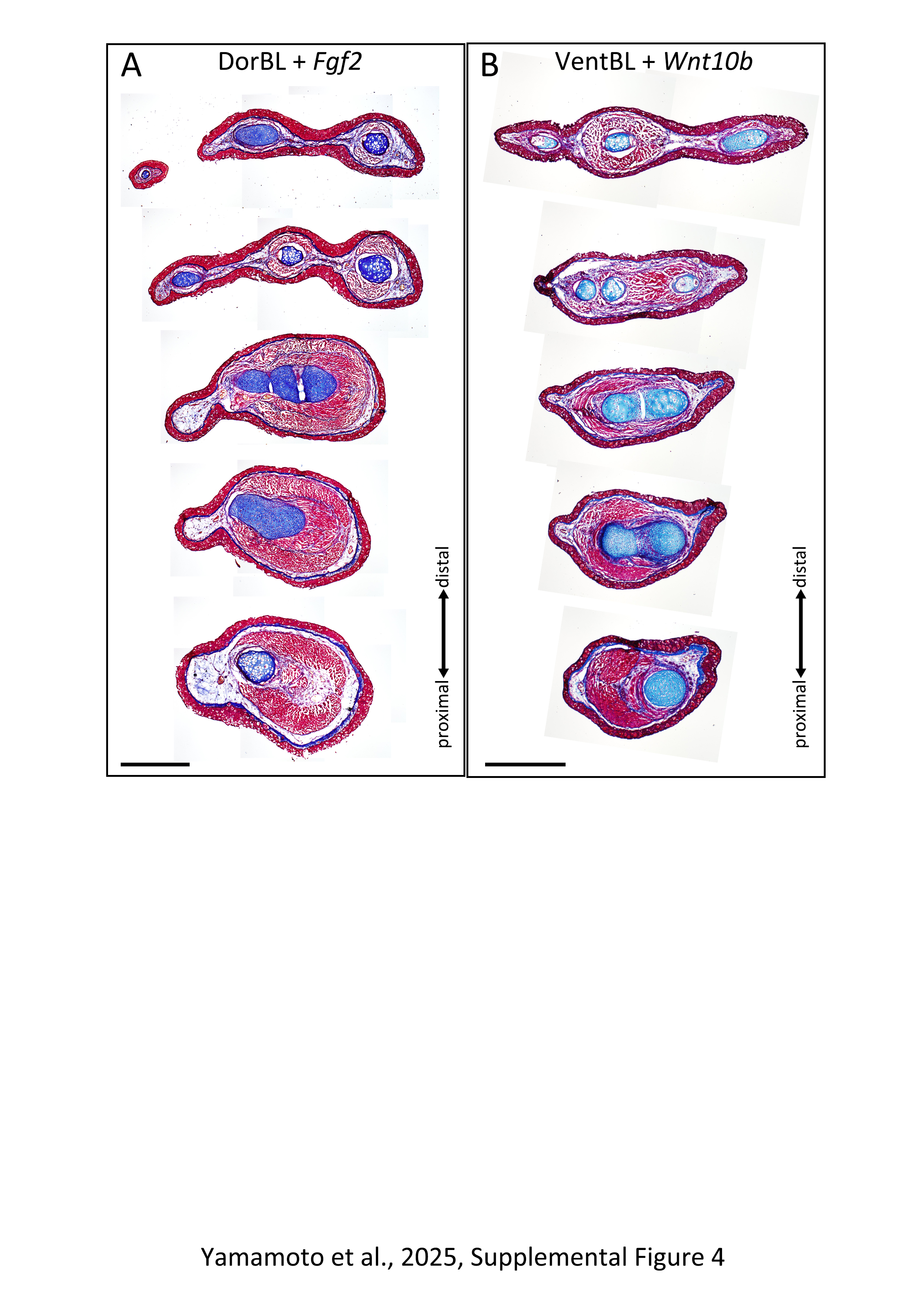

### Sup. Fig. 5

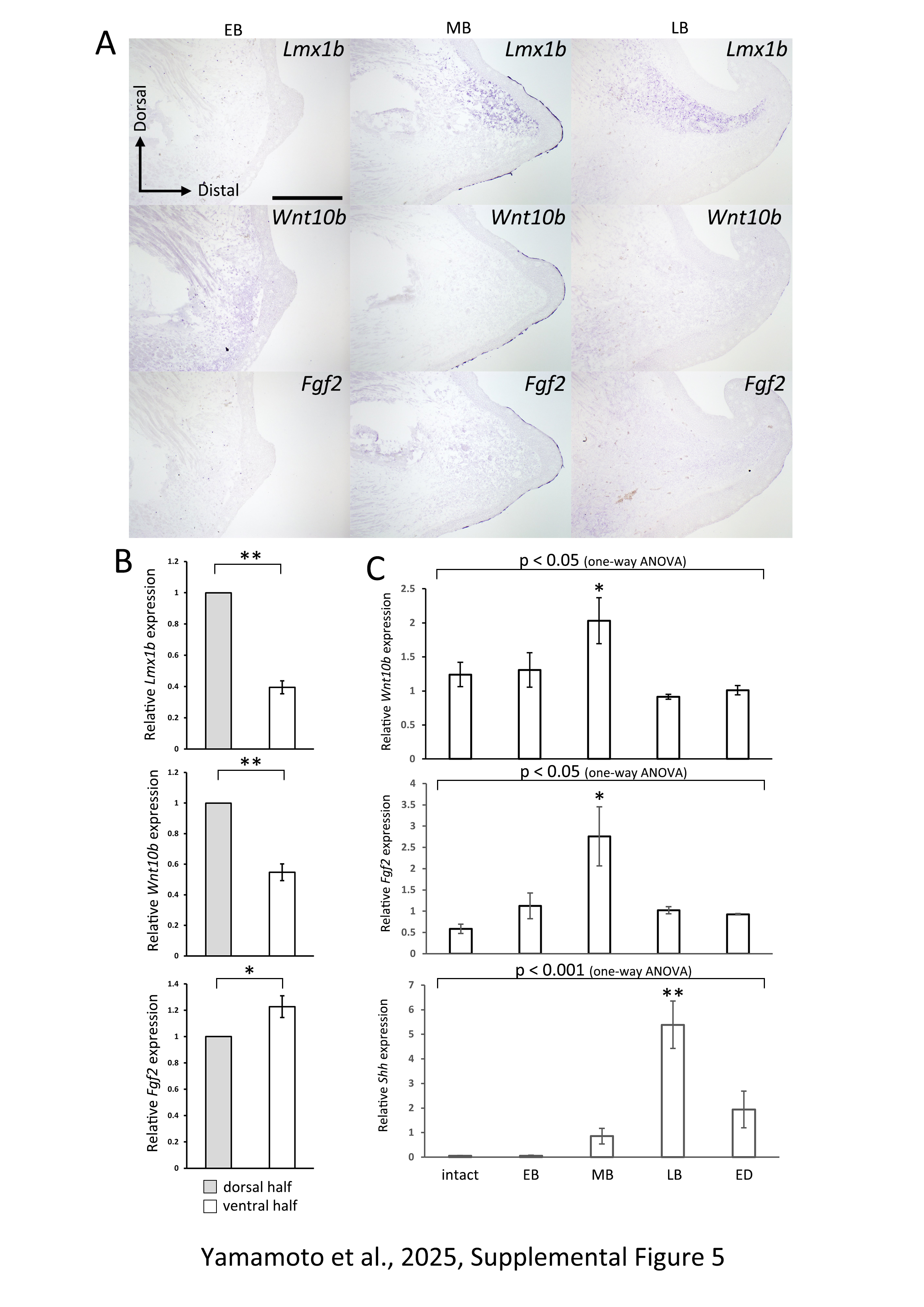

### Sup. Fig. 6

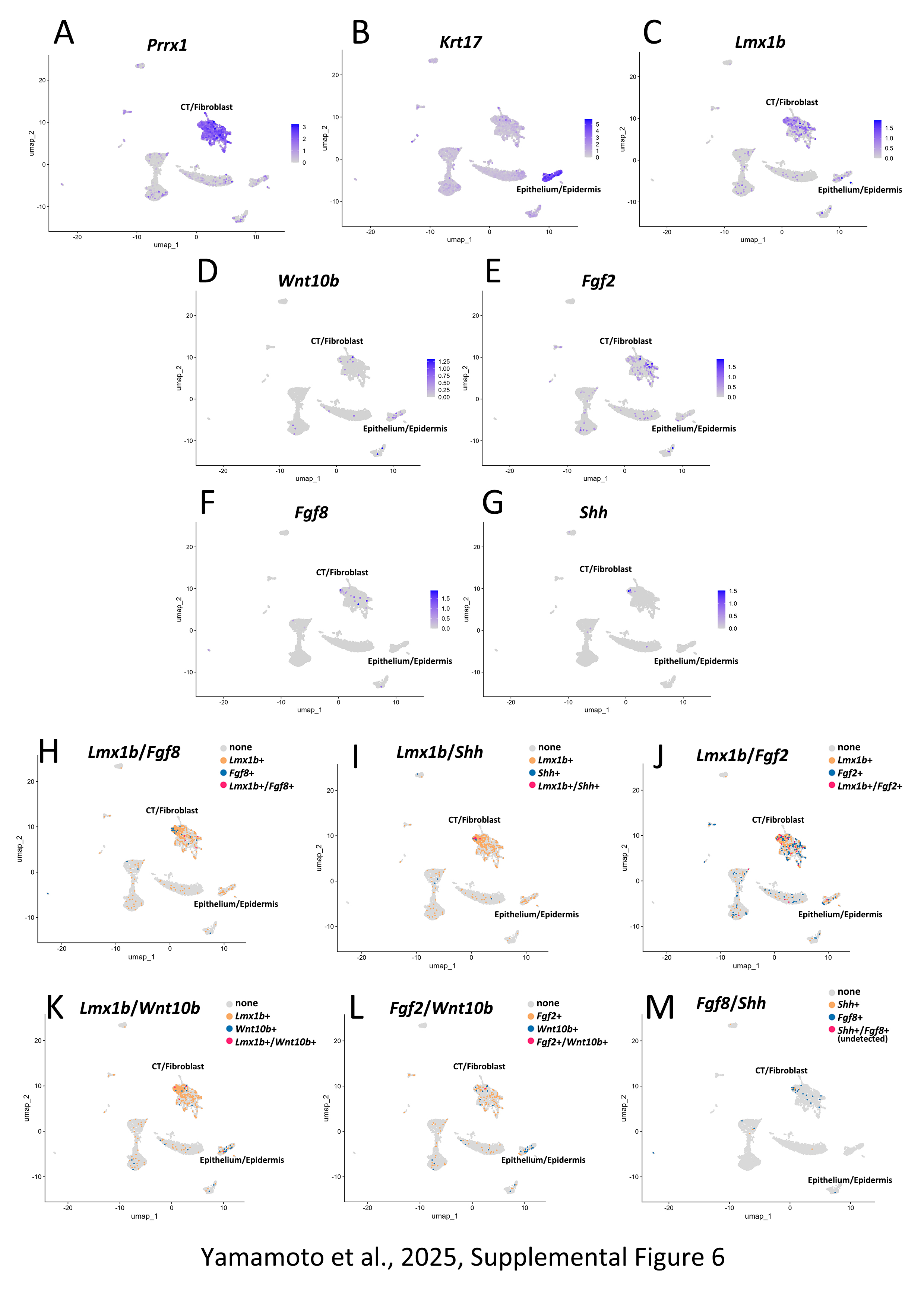
